# Colonization and diversification of aquatic insects on three Macaronesian archipelagos using 59 nuclear loci derived from a draft genome

**DOI:** 10.1101/063859

**Authors:** Sereina Rutschmann, Harald Detering, Sabrina Simon, David H. Funk, Jean-Luc Gattolliat, Samantha J. Hughes, Pedro M. Raposeiro, Rob DeSalle, Michel Sartori, Michael T Monaghan

**Affiliations:** Leibniz-Institute of Freshwater Ecology and Inland Fisheries (IGB), Müggelseedamm 301, 12587 Berlin, Germany; Berlin Center for Genomics in Biodiversity Research, Königin-Luise-Straße 6-8, 14195 Berlin, Germany; Department of Biochemistry, Genetics and Immunology, University of Vigo, 36310 Vigo, Spain; Sackler Institute for Comparative Genomics, American Museum of Natural History, Central Park West and 79^th^ St., New York, NY 10024, USA; Biosystematics Group, Wageningen University, Droevendaalsesteeg 1, 6708 PB Wageningen, The Netherlands; Stroud Water Research Center, Avondale, Pennsylvania 19311, USA; Musée cantonal de zoologie, Palais de Rumine, Place de la Riponne 6, 1014 Lausanne, Switzerland; Department of Ecology and Evolution, Biophore, University of Lausanne, 1015 Lausanne, Switzerland; Centro de Investigação e de Tecnologias Agro-Ambientais e Biológicas (CITAB), Universidade de Trás-os-Montes e Alto Douro, Quinta de Prados, Apartado 1013, 5001-801 Vila Real, Portugal; Research Centre in Biodiversity and Genetic Resources (CIBIO)-Açores and the Biology Department, University of Azores, Rua Mãe de Deus 13A, 9501-855 Ponta Delgada, Portugal

**Keywords:** Baetidae, island radiation, multispecies coalescent, phylogeny, phylogeography

## Abstract

The study of processes driving diversification requires a fully sampled and well resolved phylogeny. Multilocus approaches to the study of recent diversification provide a powerful means to study the evolutionary process, but their application remains restricted because multiple unlinked loci with suitable variation for phylogenetic or coalescent analysis are not available for most non-model taxa. Here we identify novel, putative single-copy nuclear DNA (nDNA) phylogenetic markers to study the colonization and diversification of an aquatic insect species complex, *Cloeon dipterum* L. 1761 (Ephemeroptera: Baetidae), in Macaronesia. Whole-genome sequencing data from one member of the species complex were used to identify 59 nDNA loci (32,213 base pairs), followed by Sanger sequencing of 29 individuals sampled from 13 islands of three Macaronesian archipelagos. Multispecies coalescent analyses established six putative species. Three island species formed a monophyletic clade, with one species occurring on the Azores, Europe and North America. Ancestral state reconstruction indicated at least two colonization events from the mainland (Canaries, Azores) and one within the archipelago (between Madeira and the Canaries). Random subsets of the 59 loci showed a positive linear relationship between number of loci and node support. In contrast, node support in the multispecies coalescent tree was negatively correlated with mean number of phylogenetically informative sites per locus, suggesting a complex relationship between tree resolution and marker variability. Our approach highlights the value of combining coalescent-based phylogeography, species delimitation, and phylogenetic reconstruction to resolve recent diversification events in an archipelago species complex.

## 1. Introduction

Any inference about the ecological and evolutionary processes driving diversification requires a well sampled and fully resolved phylogeny upon which traits can be mapped. Molecular phylogenetic studies historically have been limited to a small number of loci. The majority of studies are based largely on mitochondrial DNA (mtDNA) loci (Avise et al., 2000; Garrick et al., 2015) which have the benefit of small population size and high levels of polymorphism but suffer from several characteristics that can limit their suitability to reconstruct the evolutionary process. These include an inability to detect processes that confound gene trees and species trees such as hybridization and introgression, the inference of oversimplified or unresolved evolutionary relationships based on their matrilineal history, underestimated genetic diversity (Zhang and Hewitt 2003), and overestimation of divergence times (Zheng et al., 2011). Another major drawback is the presence of mtDNA genes that have been transposed to the nuclear genome, forming nuclear mitochondrial DNA (Numt; Lopez et al., 1994) which may appear homologous but give very different evolutionary signals from those of the real mtDNA. Phylogenetics has begun to benefit from more widespread use of single-copy nuclear DNA (nDNA) loci, and several recent studies have applied greater numbers of nDNA loci with success at the species (e.g. *Ambystoma tigrinum* (O’Neill et al., 2013); *Triturus cristatus* (Wielstra et al., 2014)), genus (e.g. *Takydromus* (Tseng et al., 2014); *Heliconius* (Kozak et al., 2015)), and higher taxonomic levels (e.g. Plethodontidae (Shen et al., 2016)).

The phylogenetic resolution of closely related taxa enables crucial insights in studies of evolution. In particular, the investigation of recent or ongoing species radiations helps to explain how components such as adaptation and hybridization are involved in the diversification process (e.g. Monaghan et al., 2006; Morvan et al., 2013; Giarla and Esselstyn 2015; Toussaint et al., 2015). A number of model systems in evolutionary biology come from closely related species groups that have diversified in island archipelagos (Schluter 2000; Gillespie and Roderick 2002, and references therein). Examples include Darwin’s finches (Grant and Grant 2008), *Anolis* lizards (Losos and Ricklefs 2009), or Hawaiian spiders (Gillespie et al., 1994). While a robust phylogeny is needed to study diversification and adaptation in such groups, phylogenetic analysis of close relatives can be problematic. Discordance between gene trees and species trees is more likely when speciation is recent and the effective population size of the ancestral population is large relative to the age of the species (Kubatko and Degnan 2007; Degnan et al., 2012). This discordance can arise through hybridization, gene duplication and loss, and incomplete lineage sorting (Maddison 1997; Degnan and Rosenberg 2009; Knowles and Kubatko 2010; Nakhleh 2013). Increasing arrays of methods exist for examining multilocus data that account for these processes (Rannala and Yang 2003; Edwards 2009; Heled and Drummond 2010; Knowles and Kubatko 2010). Unfortunately, the appropriate data for these analyses can be lacking because it is difficult to generate sequence data for a sufficient number of suitable nDNA loci from non-model systems. Most nDNA loci exhibit low levels of polymorphism and therefore many loci are needed, whereas identification of novel nDNA loci that are suitable as phylogenetic markers is generally not straightforward. Here we use a whole-genome draft of a non-model species to develop nDNA markers suitable for phylogenetic reconstruction.

Macaronesia consists of four archipelagos (Azores, Madeira, Canary Islands, and Cape Verde) whose flora and fauna have been used in several studies as model systems for evolutionary research. Their distances to the adjacent continental mainland vary from 110 km (Fuerteventura in Canary Islands to Morocco) to more than 2000 km (Flores in the Azores to Portugal). Several colonization pathways have been identified (Juan et al., 2000; Emerson 2002; Emerson and Kolm 2005), including a single colonization event followed by stepping-stone dispersal (Juan et al., 1997; Emerson and Oromi 2005; Illera et al., 2007; Arnedo et al., 2008; Dimitrov et al., 2008), or multiple independent colonization events within the Canary Islands (Nogales et al., 1998; Ribera et al., 2003a; Díaz-Pérez et al., 2012; Rutschmann et al., 2014; Gohli et al., 2015; Stervander et al., 2015; Faria et al., 2016). While much research has been carried out on island evolution and endemism of terrestrial organisms, comparatively limited information exists for aquatic invertebrates (e.g. Stauder 1995; Drotz 2003; Ribera et al., 2003b, 2003c; Jordal and Hewitt 2004; Hughes and Malmqvist 2005). This is a large discrepancy considering that aquatic insects contribute a disproportionally large amount of global biodiversity despite the relatively small extent of their habitat (Dijkstra et al., 2014).

Mayflies are well suited for phylogeographic studies considering their ancient origins (300 million years ago (Ma)), global distribution, and limited dispersal ability due to the strict water habitat fidelity of larvae and very short life of the winged adults (Monaghan et al., 2005; Barber-James et al., 2008). Several studies have pointed out their unusual potential for dispersion, reporting mayfly species on remote islands such as the Azores (Brinck and Scherer 1961; Raposeiro et al., 2012), trans-oceanic dispersal between Madagascar and continental Africa (Monaghan et al., 2005; Vuataz et al., 2013), and recent colonization processes of several lineages on the Canary Islands and Madeira ≈ 14 Ma, including a close link to the African mainland (Rutschmann et al., 2014).

The species complex of *Cloeon dipterum* L. 1761 is one of the most common and abundant species of freshwater insects in European standing water. The taxonomic classification and phylogenetic relationships within the *C. dipterum* s.l. species complex, including its complicated synonymy, remain largely unknown. The species complex belongs to the subgenus *Cloeon* Leach, 1815. In Europe, *Cloeon* consists of *C. dipterum*, two other currently recognized species (*C. peregrinator* Gattolliat and Sartori, 2008, and *C. saharense* Soldán and Thomas, 1983), and three species with unclear status (*species inquirenda; C. cognatum* Stephens, 1836, *C. inscriptum* Bengtsson, 1914, and *C. rabaudi* Verrier, 1949) that are often considered to be synonyms of *C. dipterum*. Its distribution ranges from North America, across Europe to Northern Asia (excluding China), making it one of the largest known distributions among mayflies (Bauernfeind and Soldán 2012, and references therein). Larvae are found in a variety of aquatic habitats, including natural standing or slow-flowing waters, brackish water, intermittent watercourses, and artificial biotopes across a wide range of climatic zones (Bauernfeind and Soldán 2012, and references therein).

For this study we used a draft genome sequence of *Cloeon* to develop 59 nDNA loci suitable for phylogenetic reconstruction of closely related members of the *C. dipterum* s.l. species complex of mayflies. We identified target genes and designed primer pairs for them. Standard PCR and Sanger sequencing were used to generate sequences. We then applied Bayesian phylogenetic inference using concatenated sequence alignments and multispecies-coalescent approaches to delineate species, examine their colonization from the mainland, and understand their diversification throughout Atlantic oceanic islands (Fig. 1, Azores, Madeira, and Canary Islands). Additionally, we quantitatively examined the effect of increasing numbers of nDNA loci on tree resolution. Our analyses show how marker development can proceed efficiently from draft whole genomes and that large numbers of nDNA loci can produce fully resolved trees in closely related taxa, revealing the evolution and diversification of the geographically widespread *C. dipterum* s.l. species complex. The disentangled colonization routes of the three species occurring on the Macaronesian Islands highlight transoceanic dispersal abilities of aquatic insects as an important driver of allopatric speciation, including sympatric occurring sister-species on the islands and the mainland.

**Fig. 1.**
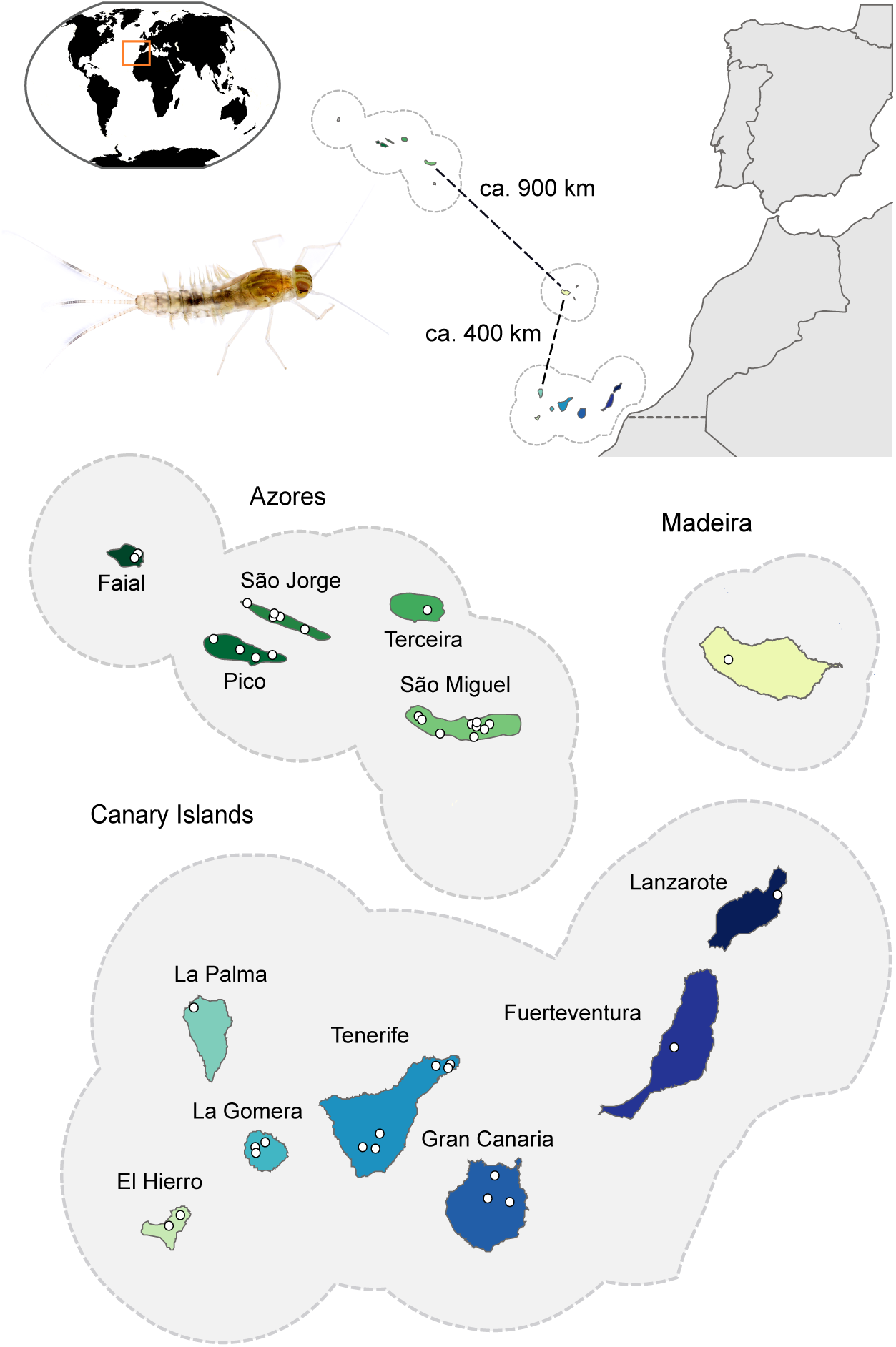
Overview of the sampling localities in the Macaronesian region. The three archipelagos of the Azores, Madeira, and the Canary Islands are shown in detail, whereby the 38 sampling sites are indicated by white dots. For the Azores and Madeira only islands with sampling sites are shown in the detailed view. Photo of *Cloeon dipterum* s.l. larvae by Amanda44/CC BY 3.0.

## 2. Material and methods

### 2.1 Development of nuclear DNA loci

To develop a set of nuclear loci we sequenced a newly created whole-genome library of *C. dipterum* (described in Rutschmann et al., 2016). Libraries were generated from laboratory-reared subimagos of *C. dipterum* specimens (full siblings). DNA was extracted from pooled specimens (five to 20) after removing eyes and wings using the Invisorb^®^ Spin Tissue Mini kit (STRATEC, Berlin, Germany). Extracted DNA was precipitated using Isopropanol and pooled to obtain higher DNA yield. We prepared one 454 shotgun and one 454 paired-end library according to the manufacturer´s guidelines (Rapid Library Preparation Method Manual, GS FLX+ Series - XL+, May 2011; Paired End Library Preparation Method Manual −20 kb and 8 kb Span, GS FLX Titanium Series, October 2009). Fragments were amplified with an emulsion PCR (emPCR Method Manual - Lib-L SV, GS FLX Titanum Series, October 2009; Rev. Jan 2010). Four lanes per library were sequenced on a Roche (454) GS FLX machine. The sequence reads were trimmed and *de novo* assembled using NEWBLER v. 2.5.3 (454 Life Sciences Corporation) with default settings for large datasets. We generated two different assemblies, one using the reads from the shotgun library and one using the reads from both shotgun and paired-end libraries. For ortholog prediction, the newly assembled draft whole genomes were combined with 4,197 expressed sequence tag (EST) sequences from *Baetis* sp. (FN198828–FN203024; Simon et al., 2009). *Cloeon* and *Baetis* belong to the Baetidae subfamilies Cloeoninae and Baetinae. Primer pairs were designed in the conserved regions of orthologous sequences from included taxa. The above analysis procedures have been incorporated into the DISCOMARK pipeline for marker discovery and primer design (Rutschmann et al., 2016; see Supplementary File 2).

### 2.2 Taxon sampling and DNA extraction

We sampled individuals of the *C. dipterum* s.l. species complex from larval aquatic habitats at 38 sampling sites on 13 islands including the Azorean archipelago, the Canary Islands, Madeira (Fig. 1), and 32 sampling sites on the European and North American mainland (Supplementary Tables 1 and 2). All samples were preserved in 99% ethanol in the field and stored at 4°C until analysis. DNA was extracted from 107 individuals using NucleoSpin^®^ 96 tissue kits (Macherey-Nagel, Düren, Germany). Our analysis included multiple populations of all currently recognized taxa (based on both morphological and molecular data) on the islands (Brinck and Scherer 1961; Gattolliat et al., 2008; Rutschmann et al., 2014).

### 2.3 PCR amplification, sequence alignment, and sequence heterogeneity

We sequenced 60 loci for the study: the mtDNA barcoding gene (*cox1*) and 59 newly developed nDNA markers from protein-coding gene regions. The *cox1* locus was amplified and sequenced using the procedure described by Rutschmann et al., 2014. Based on a general mixed Yule-coalescent (gmyc) model analysis (Fujisawa and Barraclough 2013) of *cox1*, we selected a representative set of 29 individuals, including a set of one to nine individuals for each gmyc species, depending on the number of locations at which a given gmyc species was found (see 2.4 Species assignment and population structure analysis). For these individuals, we obtained nDNA sequences using the 59 newly designed primer pairs (Supplementary Table 3). Nuclear loci were amplified using standard polymerase chain reaction (PCR) protocols with an annealing temperature of 55°C. The PCR products were custom purified and sequenced at Beckman Coulter Genomics (Essex, UK) or Macrogen (Amsterdam, The Netherlands). Forward and reverse sequences were assembled and edited using GENEIOUS R7 v.7.1.3 (Biomatters Ltd.). Length variation (i.e. heterozygous indels) was decoded using CODONCODE ALIGNER v.3.5.6 (CodonCode Corporation, Centerville MA, USA). Additionally, we included published sequences of six nDNA loci from four individuals (KU971838-KU971840, KU971851, KU971919-KU971921, KU971933, KU972490-KU972492, KU972503, KU972568-KU972570, KU972583, KU972653-KU972654, KU972666, KU973060-KU973062, KU973074; Rutschmann et al., 2016).

Multiple sequence alignments were made for each locus using MAFFT v.7.050b (L-INS-I algorithm with default settings; Katoh and Standley 2013). The predicted orthologous sequences of *Baetis* sp. were used to check the correct exon-intron splicing boundaries (canonical and non-canonical splice site pairs) of each alignment. Exon-intron boundaries of locus 411912 could not be fully reconstructed and thus we used the exon sequence predicted from tblastx searches for subsequent analyses. Locus alignments were split into coding and non-coding parts using a custom script (https://github.com/srutschmann/python_scripts/blob/master/extract_introns.py). All coding alignments were checked for indels and stop codons using Mesquite v.2.75 (Maddison and Maddison 2011). Genotypes of the coding alignments were phased using the probabilistic Bayesian algorithm implemented in PHASE v.2.1.1 (Stephens et al., 2001; Stephens and Donnelly 2003) with a cutoff value of 0.6 (Harrigan et al., 2008; Garrick et al., 2010). Multiple runs were performed for each alignment and phase calls checked for consistency. Input and output files were formatted using the scripts from SeqPHASE (Flot 2010). Heterozygous sites that could not be resolved were coded using ambiguity codes and remained in the data set for subsequent sequence analyses. All alignments were re-aligned after phasing with MAFFT. As an outgroup we used *Baetis* sp.. The number of variable sites, informative sites, and Tajima´s D for each locus were determined using a custon script and the package DENDROPY (Sukumaran and Holder 2010; https://github.com/srutschmann/python_scripts/blob/master/alignment_stats.py) and a custom script.

### 2.4 Species assignment and population structure analysis

Most analyses that use phylogenetic or multilocus species tree approaches require *a priori* species assignment. Because of the partly unknown and largely incomplete taxonomy of the group, we used two approaches to first assign the 29 *C. dipterum* individuals to putative species. The first was a gmyc analysis (Fujisawa and Barraclough 2013) of mitochondrial *coxI* and the second was a Bayesian clustering algorithm using all nuclear loci to assign individuals to ‘populations’ (STRUCTURE, Pritchard et al., 2000; Falush et al., 2003). The gmyc approach was carried out using *cox1* from 147 specimens that included all newly sequenced *Cloeon* individuals, published sequences that were available as of February 2016 (Supplementary Table 2), six newly sequenced individuals of *C. simile* Eaton, 1870, and *Baetis rhodani* (KF438126) as an outgroup. The analysis followed that of Rutschmann et al., (2014) except that we used BEAST v.2.3.2 (Bouckaert et al., 2014) and the first two codon positions were modeled with HKY + I while the third codon position was modeled with HKY + Γ. For the Bayesian clustering we used the exon_all_data matrix (see 2.5 Phylogenetic and species tree reconstructions). We assumed 1-10 genotypic clusters (K) and ran nine replicate analyses for each K, using 10^6^ MCMC generations with a burn-in of 10%. All individuals were assigned probabilistically without *a priori* knowledge to genetic clusters. We applied an admixture model with default settings (Supplementary File 3).

### 2.5 Phylogenetic and species tree reconstructions

We prepared three data matrices (Table 1), contianing all nDNA sequences (all_data), all coding genotypes (exon_all_data), and all coding haplotypes (exonhap_all_data). Because data matrices were not 100% complete (see 3.1 Development of nuclear DNA loci) we compiled a second set of matrices that were 100% complete using only the 17 loci that were successfully sequenced in all 29 individuals (complete_matrix, exon_complete_matrix, exonhap_complete_matrix). All individual locus alignments were concatenated using a Python script (https://github.com/srutschmann/python_scripts/blob/master/fasta_concat.py). We used two partitioning schemes for the phylogenetic analysis. One used the most appropriate substitution model for each locus (“partition_locus”) according to a Bayesian Information Criterion using the program jModelTest v.2.1.5 (Guindon and Gascuel 2003; Darriba et al., 2012) (Supplementary Table 3). The other used the best-fit partitioning scheme identified by PartitionFinder v.2 (https://github.com/brettc/partitionfinder) (“partition_whole”). The latter was determined using the greedy algorithm (Lanfear et al., 2012), whereby PhyML (Guindon et al., 2010) was used for model evaluation prior to the Bayesian analysis, and RAxML v.8.2.8 (Stamatakis 2014) was used for model evaluation prior to maximum likelihood analysis (see 2.7 Relationship between node support and number of loci).

**Table 1.**
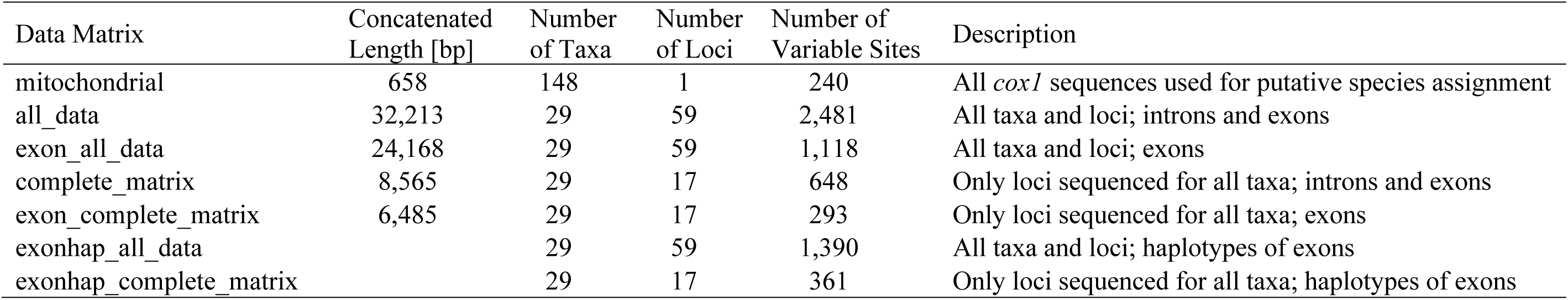
Overview of seven sequence alignments, including one based on mitochondrial sequences and six concatenated matrices based on nuclear sequences. The mitochondrial sequence alignment including only the species group of *Cloeon dipterum* s.l. comprised 130 taxa with 148 variable sites. Matrices containing all taxa and all loci were >75% complete; the matrices containing only loci sequenced for all taxa were 100% complete. Exon matrices refer to exon sequence alignments and the exonhap matrices refer to exon haplotype sequences.

Bayesian phylogenetic analysis was carried out on each partitioning scheme using MRBAYES v.3.2.3 (Ronquist et al., 2012) using exons, one analysis using all data and one using only the complete matrix (Table 1; exon_all_data, exon_complete_matrix). We unlinked the nucleotide frequencies, gamma distributions, substitution rates and the proportion of invariant sites across partitions. Each run consisted of two independent analyses of four MCMC chains, each with 10^7^ generations and 25% burn-in.

Species tree reconstructions were carried out under a multispecies coalescent framework (Drummond and Rambaut 2007; Heled and Drummond 2010) as implemented in the program *BEAST v.2.1.3 (Bouckaert et al., 2014). All analyses were performed using exons, one analysis using all data and one using only the complete matrix (Table 1; exonhap_all_data, exonhap_complete_matrix). All individuals were *a priori* assigned to species based on the gmyc and Bayesian clustering analyses (see 2.4 Species assignment and population structure analysis). In the Bayesian clustering analysis, one individual sampled from Russia was considered to be admixed based on Bayesian posterior probability (PP) assignment values > 0.05 for more than one cluster (Supplementary File 3, Supplementary Fig. 1, Supplementary Table 5). This individual was therefore excluded from further analysis. We used a relaxed uncorrelated lognormal clock for gene tree estimation at each locus and a Yule speciation-process prior. Six independent runs of 8 × 10^8^ million generations each were conducted. Runs were combined in LOGCOMBINER v.2.1.3 (Bouckaert et al., 2014), whereby all parameters reached effective sample sizes (ESS) > 600. Maximum clade credibility trees for each species tree were obtained using TREEANNOTATOR v.2.1.3 (Bouckaert et al., 2014).

### 2.6 Ancestral state reconstruction

Ancestral state reconstruction was used to test the geographical direction of the radiation (i.e. Continental to Island or Island to Continental). Ancestral range patterns of each individual were defined into four geographic areas: (1) a broadly defined Continental referring to the European and North American mainland, (2) Canary Islands, (3) Madeira, and (4) Azores. As input tree, we used the concatenated tree based on the exon_all_data inferred with MRBAYES. A chronogram was fit to the tree using the chronos function in the ape v.3.4 (Paradis et al., 2004) package for R. Ancestral states were estimated under an equal-rates (ER) model using the function ace, and the scaled likelihoods of each ancestral state were calculated using the function lik.anc. An MCMC approach was used to sample character histories from their PP distribution generating 1,000 stochastic character maps with the function make.simmap of the phytools v.0.4.98 (Revell 2012) package in R (R Core Team, 2016; Supplementary File 4).

### 2.7 Relationship between node support and number of loci

To investigate how the number of analyzed loci affected node support values, we performed phylogenetic reconstructions based on multiple subsets of randomly selected loci (Supplementary Table 4). This included twelve subsets for the phylogenetic analysis and four subsets for the multispecies coalescent analysis. For the Bayesian phylogenetic analysis of subsets of loci we used MRBAYES (see 2.5 Phylogenetic and species tree reconstructions), applying both partition schemes. As an alternative to Bayesian PP, we calculated Shimodaira-Hasegawa approximate likelihood ratio test (SH-aLRT; Guindon et al., 2010) supports of each node using maximum likelihood phylogenies. For this, we first estimated the best tree with RAXML v.8.2.9 applying the “partition_whole” partitioning scheme (Supplementary Table 4) and rapid bootstrap analysis with 1,000 replicates. Linear regressions were used to model the number of supported nodes for PP ≥ 0.95 and PP = 1, and the SH-aLRT supports ≥ 0.95 and SH-aLRT supports = 1 as a function of the number of loci used in the analysis. The Pearson correlation between the number of loci and number of supported nodes was calculated using the stats package for R.

The multispecies coalescent tree reconstructions were performed and summarized as described in section 2.5. We calculated the correlation between the mean number of parsimony-informative sites per locus and the mean node support values of the randomly selected loci and the resulting tree reconstruction using the cor.test function of the stats package for R.

## 3. Results

### 3.1 Development of nuclear DNA loci

Whole-genome sequencing resulted in 1,109,684 raw reads, including 651,306 reads for the shotgun library and 458,378 reads for the paired-end library, with an average large contig length of 1,187 and 736 base pairs (bp), respectively (BioSample SAMN03202660, BioProject PRJNA268073, Sequence Read Archive SRP050093). All reads were assembled into 68,473 contigs with an N50 of 1,116 bp. The reads of the shotgun library were assembled into 31,827 contigs with an N50 of 1,260 bp. We detected 918 putative orthologous gene sequences for *C. dipterum* from the contigs derived from the shotgun library, 1,298 putative orthologous gene sequences from the contigs of the combined assembly, and 416 for *Baetis* sp. (Supplementary Table 6). From these, we haphazardly selected 65 markers for primer design, approximately 80% of which included orthologous sequences from both taxa. These were chosen based on the presence of conserved regions and short introns suitable for primer design. Based on preliminary laboratory testing, 59 markers were selected that amplified consistently and had similar annealing temperatures in order to simplify the large number of PCR reactions (Supplementary Table 3).

Total fragment length per sequenced locus ranged from 210 - 1,007 bp with a mean of 545 bp. Exon sequence length ranged from 210 - 710 bp with a mean of 410 bp (Supplementary Table 3) (KF438124-KF438125, KU757080-KU757184, and KU971616-KU973191). The full data matrix of all 29 individuals and all 59 loci including exons and introns (all_data) was 32,213 bp in length when concatenated and when introns were removed (exon_all_data) it was 24,168 bp (Table 1). All individuals were successfully sequenced for at least 44 loci, and the above matrices were >75% complete. The 100% complete matrix included 17 loci that were sequenced successfully for all 29 individuals. All heterozygous indels were located in the intron sequences. However, 100 heterozygous sites could not been resolved and remained in the exonhap alignments.

The number of variable sites per locus ranged from six to 65 (mean: 18.95). In the exon_all_data matrix, there was one SNP per every 21.62 nucleotides sequenced (i.e. total length per total number of variable sites). The loci included between six and 54 informative sites (mean: 16) and one to 26 ambiguous sites (mean: 8.4). Nucleotide diversity ranged from 0.007 to 0.04 (mean: 0.017), and Tajima´s D varied between −0.85 and 1.97 (mean: 0.29) (Supplementary Table 7).

### 3.2 Species assignment and population structure

There were 62 unique *cox1* haplotypes of *Cloeon* and the gmyc model was a significantly better fit to the data than the null model (χ^2^ = 31.00, *p* < 0.001). Seven putative species were delineated within *C. dipterum* s.l. (Fig. 2a): One occurred only in Asia (South Korea) while the remaining six comprised three species with distributions that included the Macaronesian Islands (IS1, IS2, IS3) and three species only occurring on the European and North American continents (CT1, CT2, CT3). The population assignments from the Bayesian clustering analyses of nDNA agreed with the results from the gmyc analysis for the six species of interest (Supplementary File 3, Supplementary Fig. 1, and Supplementary Table 5). No nuclear data were available for the seventh gmyc species from Asia so it was not included in the clustering analysis. Among the six, one widespread species (IS1) was found on all Azorean islands, in Greece and Italy, and in North America; one was found on the Canary Islands and Madeira (IS2), and one was found only on four Canarian islands (IS3). The gmyc model delineated all seven *C. dipterum* species even when using the most conservative estimate (95% confidence interval based on two log likelihood units: 16-19 gmyc species). The two *C. cognatum* specimens from the North American DNA barcoding project (Webb et al., 2012) had *cox1* haplotypes identical to our gmyc species IS1.

**Fig. 2.**
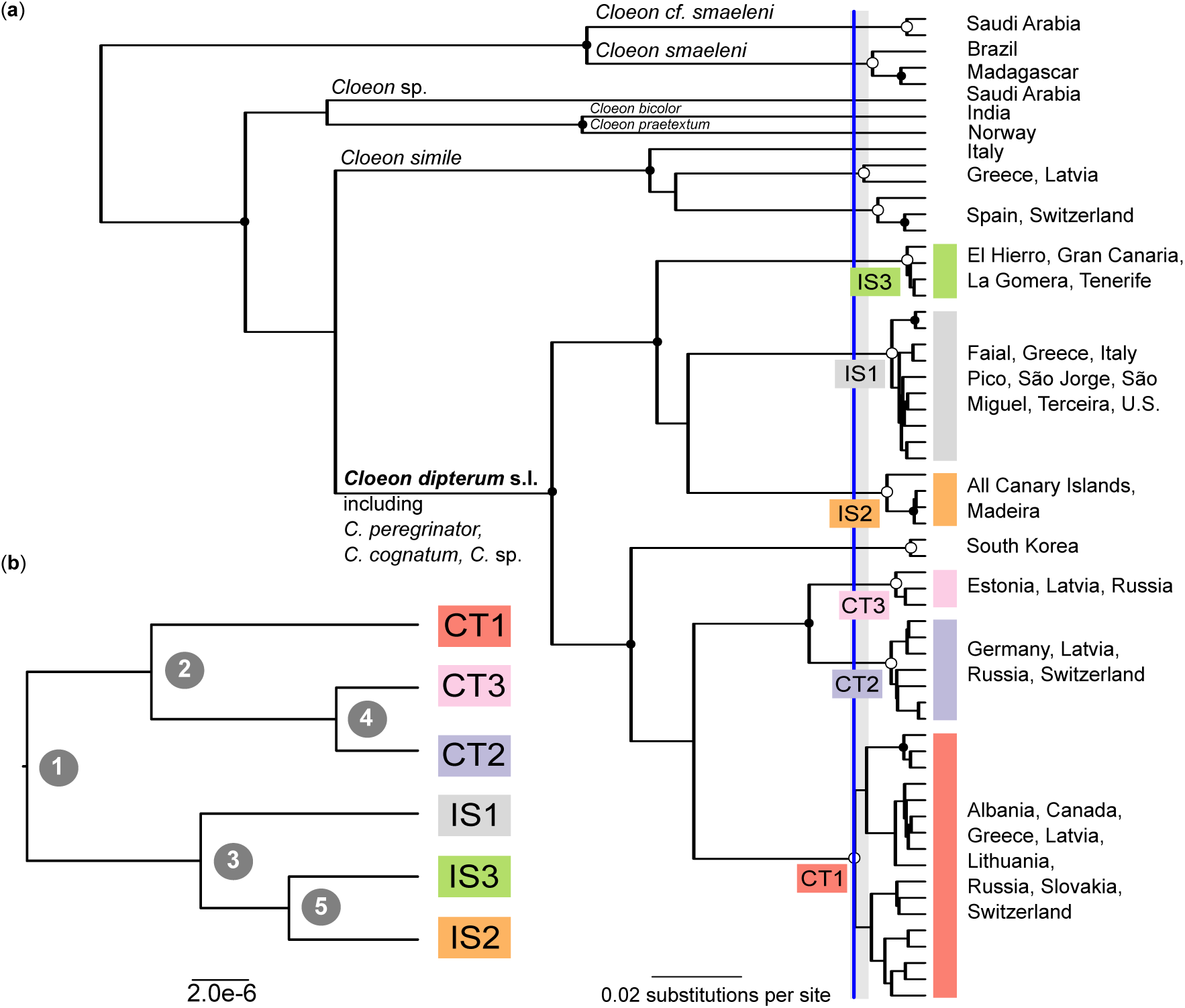
Mitochondrial gene tree and nuclear species tree topology. Species delimitation based on (**a**) the general mixed Yule-coalescent (gmyc) approach using mitochondrial data (single-locus); and (**b**) the multispecies coalescent approach using nuclear data (59 loci). An ultrametric mitochondrial *cox1* gene tree was used as input for the gmyc analysis of *Cloeon* sp (**a**). All circles indicate well supported nodes (Bayesian posterior probability ≥ 0.95). Open circles at subtending nodes indicate sequence clusters corresponding to single gmyc species. Colored rectangles and alphanumeric codes indicate the six gmyc species within *C. dipterum* s.l. The outgroup *Baetis rhodani* is not shown. The vertical blue line indicates the point of maximum-likelihood fit of the single-threshold gmyc model and 95% confidence intervals are indicated by grey shading. Terminal labels indicate sampling regions (Supplementary Table 1). The species trees of *C. dipterum* s.l. inferred using a multispecies coalescent approach based on the exonhap_all_data matrix (**b**). Posterior probabilities of the five nodes varied with the number of loci analysed and these are indicated in Table 2.

**Table 2.** Node support values of the species tree analysis (Fig. 2b) using six different sets of loci (Supplementary Table 4). Support values are given as Bayesian posterior probability (PP).

| Number<br>of Loci | Node Support |  |  |  |  | Mean |
| --- | --- | --- | --- | --- | --- | --- |
|  | All<br>(1) | Continental<br>(2) | Island<br>(3) | CT2+CT3<br>(4) | IS2+IS3<br>(5) |  |
| 17 | 1.00 | 0.99 | 1.00 | 1.00 | 1.00 | 1.00 |
| 20 | 1.00 | 1.00 | - | 1.00 | - | 0.60 |
| 30 | 1.00 | 0.85 | 0.87 | 0.87 | 0.99 | 0.92 |
| 40 | 1.00 | 1.00 | 1.00 | 0.67 | 0.83 | 0.70 |
| 50 | 1.00 | 0.83 | 0.83 | 0.83 | 1.00 | 0.87 |
| 59 | 1.00 | 0.83 | 0.83 | 0.83 | 1.00 | 0.87 |

### 3.3 Phylogenetic reconstruction

Analyses based on both exon matrices (exon_all_data; exon_complete_matrix) and both partition schemes recovered the same tree topology with strong node support, resolving each of the three species occurring on Macaronesia (IS1-IS3) as monophyletic and members of a monophyletic ‘Island clade’ (Fig. 3). The geographically widespread species IS1 was sister taxon to the two others. Species CT2 and CT3 were both monophyletic and sister group to the Island clade (Fig. 3). All individuals in CT1 were monophyletic except for a single individual that was sister taxon to the entire *C. dipterum* s.l. lineage. There were 27 resolved (PP ≥ 0.95) nodes in the tree resulting from the large, >75% complete matrix with all available data (exon_all_data, 59 loci). The only unresolved node was between the two Azorean individuals (Fig. 3). In contrast, the tree resulting from the smaller matrix with no missing data (exon_complete_matrix, 17 loci) contained only 19 resolved nodes, with lack of resolution most pronounced in IS2 (Supplementary Fig. 2).

**Fig. 3.**
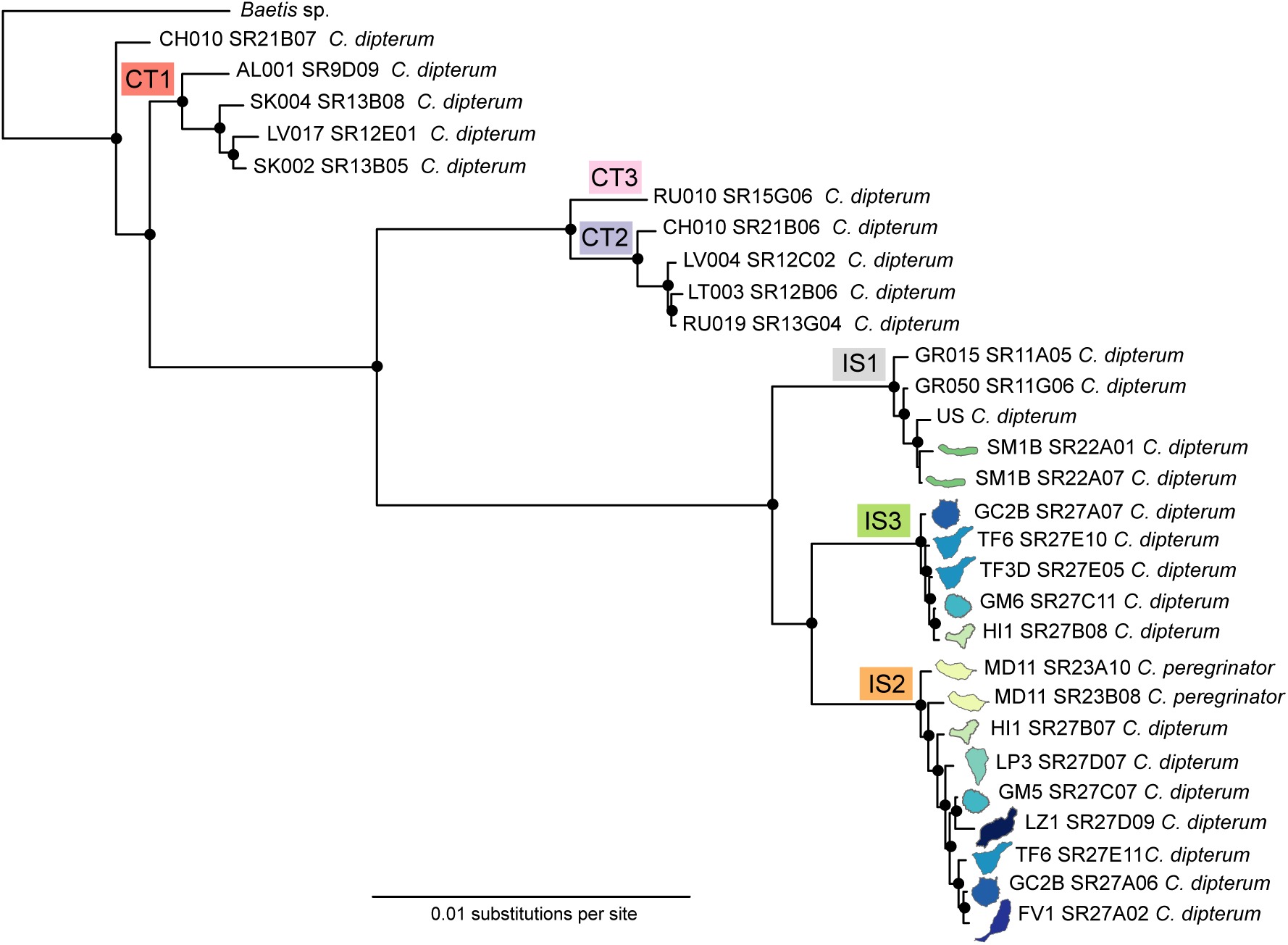
Phylogenetic relationships among *Cloeon dipterum* s.l. (including *C. peregrinator*) based on the Bayesian reconstruction using the concatenated exon_all_data matrix of 59 nuclear and one mitochondrial marker. Filled circles indicate nodes with Bayesian posterior probability ≥ 0.95. Species inferred from multilocus coalescent and gmyc analyses are indicated with colored rectangles and alphanumeric codes as in Fig. 2.

All species tree phylogenies had identical topologies and these matched the Bayesian phylogenies (Figs. 2b, 3a) in that Island and Continental clades were both monophyletic, with IS1 sister taxon to IS2 + IS3, and with CT1 sister taxon to CT2 + CT3. Using the exon_complete_matrix, all nodes were highly supported (PP ≥ 0.99; Table 2). All individuals clustered into six species in the same way in both the multilocus nDNA tree and the single-locus (*cox1*) mtDNA tree (PP = 1), but the relationships among the species were different. The mtDNA tree did not support the sister relationship of IS2 + IS3 or the monophyly of the Continental clade (Fig. 2a; Table 2).

The analysis of subsets of loci showed a positive relationship between the number of loci employed and the number of nodes resolved for phylogenetic analyses (both Bayesian and maximum likelihood) (Fig. 4). The results for the multispecies coalescent were not as clear. Support of key nodes varied widely with the number of loci employed (Fig. 2b, Table 2). The highest overall support came from analysis of 17 and 40 loci, although only the analysis using 20 loci failed to recover either node in the Macaronesian clade and resulted in no resolution other than continental monophyly (Fig. 2b, Table 2). There was a strong negative correlation (Pearson R = −0.95, *p* < 0.05) between the mean number of informative sites per locus and mean node support (Supplementary Tables 4, 7).

**Fig. 4.**
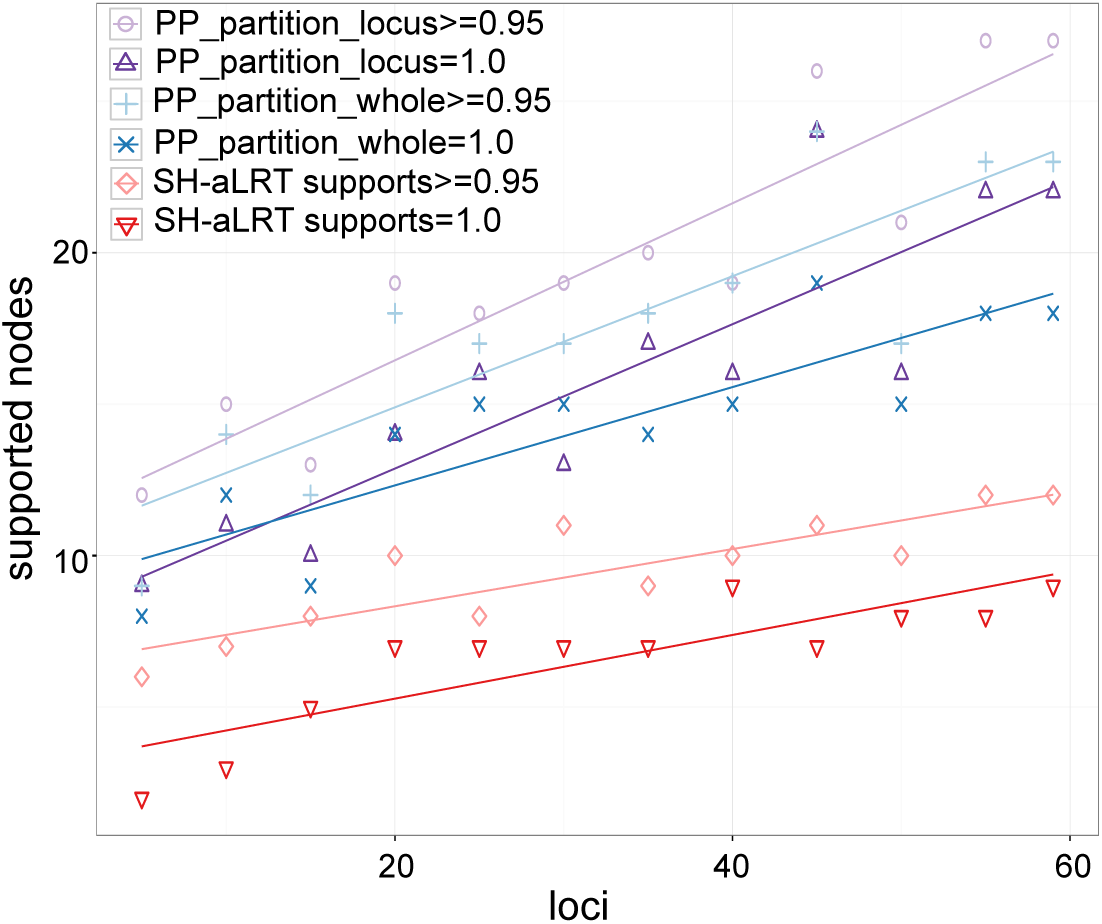
Linear relationships between number of supported nodes and number of loci analyzed, including as node support values the Posterior probability (PP) and Shimodaira-Hasegawa approximate likelihood ratio (SH-aLRT) supports for the two partition schemes, using each locus as partition (partition_locus) and using the whole alignment as one partition (partition_whole). Purple circles for PP_partition_locus ≥ 0.95 (R^2^ = 0.83, *p* < 0.001), purple triangles for PP_partition_locus = 1 (R^2^ = 0.75, *p* < 0.001), blue pluses for PP_partition_whole ≥ 0.95 (R^2^ = 0.72, *p* < 0.001), blue crosses for PP_partition_whole = 1 (R^2^ = 0.72, *p* < 0.001), red squares for SH-aLRT supports ≥ 0.95 (R^2^ = 0.74, *p* < 0.001), and red triangles for SH-aLRT supports =1 (R^2^ = 0.71, *p* < 0.001). Sets of loci are reported in Supplementary Table 4.

### 3.4 Ancestral state reconstruction

The ancestral state reconstruction identified four nodes having marginal states with less than 0.9 PP for one character, including the sister relationship between the individual CH010_SR21B07 and the remaining species, the ancestral node of IS2 + IS3 (Canary Islands and Madeira), and the nodes separating Madeiran from Canarian individuals within IS2 (Fig. 5). The Island clade had a continental origin, further a Canarian origin was estimated for IS2 + IS3. The clade IS2 was estimated to have an ancestral state of 0.59 for Madeira and 0.4 for the Canary Islands (Supplementary File 4).

**Fig. 5.**
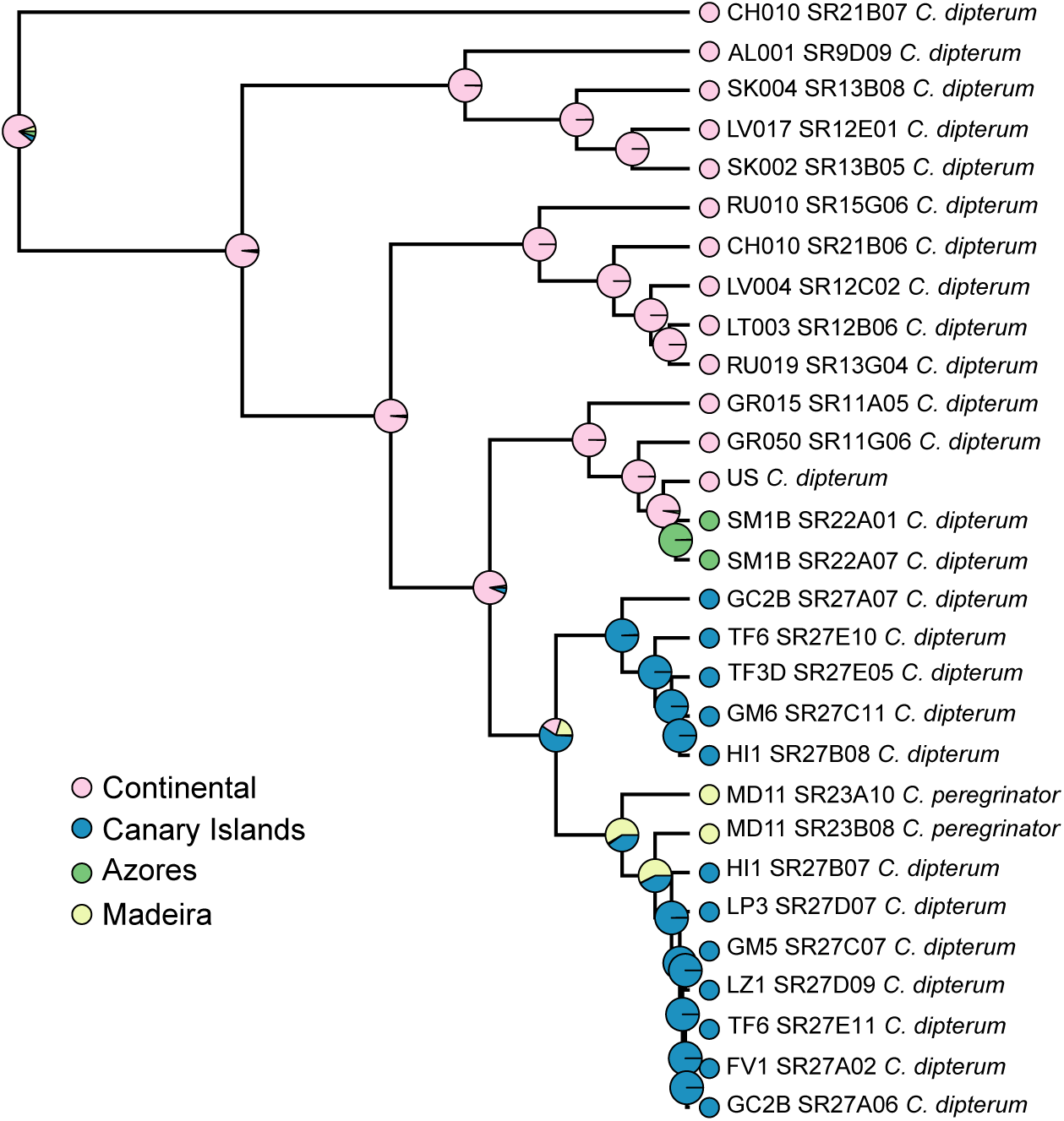
Ancestral state reconstruction of the origin estimated under an equal-rates model based on the concatenated tree. Colors highlight the four defined origins : Continental, including European and North American mainland (pink), Azores (green), Madeira (yellow), and Canary Islands (blue).

## 4. Discussion

### 4.1 Species delineation

The agreement of the multispecies coalescent and mitochondrial gmyc approaches for *a priori* species delineation support the use of *cox1* as barcoding gene for the taxa studied (e.g. Lucentini et al., 2011; Pereira-da-Conceicoa et al., 2012; Webb et al., 2012; Rutschmann et al., 2014; Gattolliat et al., 2015). Nonetheless, the gene tree based on *cox1* did not resolve the relationships among the six closely related *C*. *dipterum* s.l., with no support for the sister relationship of IS2 + IS3 or the monophyly of the Continental clade. The distant clustering of one individual (CH010_SR21B07) in the concatenated tree analyses might be explained by incomplete lineage sorting since the species tree inferences using *BEAST did result in a clear clustering of CT1 with low frequency of different topology (Fig. 3). Moreover, when incomplete lineage sorting is present, standard methods for estimating species trees, such as concatenation and consensus methods, can be statistically inconsistent (Degnan et al., 2009; Roch and Steel 2014), and produce highly supported but incorrect trees (Kubatko and Degnan 2007). The majority of gene trees could support an incorrect species tree if the phylogeny is in the anomaly zone (Degnan and Rosenberg 2006). Here this does not appear to be the case, because one would expect the concatenation and coalescent approach to support different topologies (Kubatko and Degnan 2007; Liu and Edwards 2009).

### 4.2 Species diversity

The use of nDNA and geographically extensive sampling uncovered a largely underestimated species diversity for *C. dipterum* s.l. species complex, supporting the existence of six geographically relevant species from our study (with a seventh in Asia). Recent evidence from the study of other mayfly species found fine-scale ecological differences among cryptic species detected with molecular methods (Leys et al., 2016; Macher et al., 2016), lending support to the ecological and evolutionary significance of these and other DNA-based findings. Another widespread species, *Baetis harrisoni*, was also found to consist of several cryptic species (Pereira-da-Conceicoa et al., 2012). In light of the unusually broad ecological tolerance (among mayflies) observed for *C. dipterum*, we also conclude that the lineage clearly consists of multiple independent species, as has been recognized by morphological taxonomy for some of the members (e.g. *C. peregrinator*). All of our analyses grouped the two specimens of *C. peregrinator* from Madeira with individuals from several Canary Islands into species IS2. Gattolliat et al., (2008) described *C. peregrinator* as an endemic Madeiran species based on morphological characters and support from mtDNA cytochrome-oxidase *b* sequences. At the time, there were no nDNA sequences of Canarian *C. dipterum* s.l. specimens available. Rutschmann et al., (2014) assigned all Madeiran *Cloeon* individuals to *C. peregrinator* for their mtDNA phylogeny, but the specimens were not included in their gmyc analysis because there were no *cox1* sequences available. Based on our findings here, there is no endemic *Cloeon* species on Madeira.

The focus of our study was Macaronesia and therefore nDNA results are only applicable to these taxa, but the mtDNA gene tree provides evidence for broad cryptic diversity within the subfamily Cloeoninae. *Cloeon simile* included two geographically widespread European gmyc species, and *C. smaeleni* Lestage 1924 was two gmyc species, one with Saudi Arabian and one with Afrotropical distribution. The species *C. praetexum* was clearly distinct from all other examined European specimens, which was surprising because it is thought to belong to *C. simile* s.l.. The two specimens of *C. cognatum*, which is thought to be a junior synonym of *C. dipterum* by some authors, were nested within the IS1 clade. All of the above findings must be considered preliminary because they are based on mtDNA, although we note that mtDNA and nDNA markers agreed in all of the *Cloeon* species that were directly compared. Further studies on these other *Cloeon* taxa with nuclear markers, using morphological characteristics, and including comparisons with previously described species that are now considered junior synonyms or *species inquirenda* would be a valuable complement to the work presented here.

### 4.3 Evolution, colonization, and diversification

For the species occurring in the Macaronesian region, one species appeared widely distributed on all Canary Islands and Madeira (IS2), one species was found only on the western group of the Canary Islands (IS3), and one species was found on five islands of the Azores, as well as in Italy, Greece and North America (IS1). The short branches of individuals from IS1 support very recent or perhaps ongoing gene flow. Other studies have found evidence for recent or ongoing dispersal in *Cloeon* (e.g., Monaghan et al., 2005) including a recent introduction of African *Cloeon* to South America (*C. smaeleni*, Salles et al., 2014). This long-distance dispersal ability is probably at least partly related to their reproductive flexibility including ovovivipary and their ability to survive in anthropogenic habitats. Our ancestral state reconstruction indicated that IS2 may have first colonized Madeira and then the Canaries from west to east. Colonization routes between these two archipelagos have been suggested for several taxa (Emerson et al., 2000a; Emerson et al., 2000b; Trusty et al., 2005; Illera et al., 2007; Dimitrov et al., 2008; Amorim et al., 2012). The species IS3 seems not to have reached La Palma and the two most eastern Canarian islands of Fuerteventura and Lanzarote. The dispersal of IS3 appears to have followed the progression rule, in which older islands are inhabited by older clades, which is further supported by stepping-stone dispersal along an east-western gradient.

Our data confirm at least three and possibly four independent colonization events of the islands studied, with a European origin for the Macaronesian *C. dipterum* s.l.. However, long branches between the Continental clades and the Island clade suggest there may be missing intermediates. These may occur in the Iberian Peninsula or North Africa. Several studies have proposed a North African origin for both the Canarian and Madeiran fauna (Brunton and Hurst 1998; Kvist et al., 2005; Weingartner et al., 2006; Gohli et al., 2015; Stervander et al., 2015). The Continental clades are also distantly related to one another and the long branches within both clades, compared to the Island clades, suggest there may be additional European species that are not included here.

Our results suggest a strong effect of different habitat preferences between the two Canarian species, which might affect their colonization success. Although our dataset was not quantitative, we observed that species IS3 generally occurred on islands with more potential habitats in comparison to IS2, which seems to have better dispersal abilities and might therefore be able to more successfully colonize islands with very little water occurrence. This pattern may be linked with the occurrence of suitable water habitats on the Canarian Islands. The four islands of Gran Canaria, Tenerife, La Gomera, and La Palma all have permanent natural water sources, and the island of El Hierro has several artificial water habitats due to the mostly temperate climatic conditions. In contrast, there are only a few habitats on Fuerteventura and Lanzarote due to the arid climatic conditions. The effect of habitat use on species richness has been shown for aquatic beetles (Ribera et al., 2003c), with running water bodies generally containing more species than standing ones. This pattern also applies to the Macaronesian mayflies. The genus *Baetis* occurs in running waters and is species-rich, including eight island endemic species on five islands of Madeira and the Canary Islands (Rutschmann et al., 2014). In contrast, the genus *Cloeon* comprises three species, none of which are restricted to a single island. The impact of agriculture and tourism on natural habitats (Malmqvist et al., 1995; Nilsson et al., 1998) has clearly threatened the occurrence of species living in running water habitats (*Baetis canariensis* and *B. pseudorhodani*, Rutschmann et al., 2014), but it may have had less of an effect on *C. dipterum*. The records of mayflies from El Hierro indicate a recent anthropogenic import of the species, moreover because it is the youngest island of the Canarian archipelago and its remote geographical position.

Interestingly, there were eight sampling sites (out of 32 examined sites, i.e. 25%) in which both species IS2 and IS3 occurred sympatrically. Four of these localities were natural habitats. However, more work needs to be done to make quantitative assessments on species occurrence and local abundance of the two distinct species occurring on the same habitats. A wider geographic sampling, focusing on specimens from the European mainland and North Africa will be needed to clarify the origin and distribution of the *C. dipterum* s.l. species complex. We expect to find more individuals from distinct geographic localities belonging to the species IS1, since this species appears to exhibit trans-oceanic dispersal abilities.

### 4.4 Number of loci for phylogenetics

A recent study by O’Neill et al., (2013) examined how multilocus species tree inferences varied with differing numbers of loci. Their analysis based on the 20 and 30 most informative loci (using a parsimony criterion) in their data set resulted in high PPs, whereas node support values were lower and likelihoods failed to converge when loci that were less informative were added to the analysis. They concluded this was the result of the increasing number of parameters while adding loci with decreasing levels of information. Our results are not directly comparable to those of O’Neill et al. (2013) for the species tree reconstruction, because we did not explicitly order loci by parsimony-informative sites in our tests. Nonetheless, we found a strong negative correlation between the mean number of informative sites per locus and mean node support in the coalescent species tree. This suggests that the number of informative sites was not able to explain variation in support alone, and that multiple characteristics of individual loci play an important role in whether or not analyses achieve convergence and tree resolution. For the species phylogeny, we found a positive linear correlation between number of loci and node support. This was despite the larger number of parameters. In our study we observed that the reduction in node support when using a reduced set of loci (exon_complete_matrix *vs.* exon_all_data, see 2.3 PCR amplification, sequence alignment, and sequence heterogeneity) primarily affected the most derived clade (IS2), which highlights the importance of large nDNA marker sets for the reconstruction of shallow phylogenies.

## 5. Conclusion

Our aims were to delineate the species boundaries within the *C. dipterum* species complex and place these lineages within a phylogenetic framework, in order to better understand their evolution on the Macaronesian Islands. Robust phylogenetic reconstruction of such closely related species can be challenging, but is a necessary step in the understanding of evolutionary processes of diversification and adaptation. Most of the widely used nDNA loci (e.g. rRNA) do not exhibit suitable polymorphism, resulting in a large dependence on mtDNA for phylogenetics (Garrick et al., 2015). A distinct advantage of using multiple nDNA loci comes from the advent of multilocus species tree reconstruction methods. These are important tools in the reconstruction of relationships between close relatives, which is often intractable based on single-locus (i.e. mtDNA) data. The difficulty in developing large numbers of nDNA loci remains one of the primary reasons that there are few model systems available for detailed studies of speciation and diversification processes. This is particularly true for freshwater insects, despite their overwhelming contribution to global biodiversity (Dijkstra et al., 2014). Here we developed a large set of nDNA loci using draft whole-genome sequencing. All of the procedures we used have since been incorporated into a single analysis pipeline (Rutschmann et al., 2016). Our results show that even for taxa with very limited available genomic resources, it is possible to develop sets of nuclear loci that produce fully resolved and supported coalescent-based species trees and species-level phylogenetic trees. Using these results, we were able to infer species boundaries within the largely cryptic *C. dipterum* s.l. species complex and reconstruct the diversification and island colonization history of these species with confidence.

## Author contributions

S.R., M.S. and M.T.M. conceived the study. S.R., H.D., D.H.F., J.-L.G., S.J.H., P.M.R., and M.S. collected and identified specimens. H.D., S.S., and R.D. contributed analytical tools. S.R. and M.T.M. performed and interpreted the analyses, and wrote the manuscript. All authors commented and approved the final manuscript.

## Acknowledgements

We are grateful to Katrin Preuß, Susan Mbedi, Berta Ortiz Crespo, and Lydia Wächter for laboratory work; Matthias F. Geiger, Katharina Kurzrock, Konstantinos C. Gritzalis, Andrey Przhiboro, Maria Alp, Vicenc Acuna, Peter Manko, Dávid Murányi, André Wagner, Tomas Ruginis, Luis F. Pires Braz, and Verena Lubini for field work and providing samples; Peter Rutschmann and the HPC Service of ZEDAT, Freie Universität Berlin for access to high-performance computing resources. We are greatly indebted to Marcos Báez for the precious help for the fieldwork on the Canary Islands, the Canarian authorities who provided us with the collection permissions: Servicio Administrativo de Medio Ambiente, Cabildo de Tenerife (reg. number 2014-00200/2014), Ministerio de Medio Ambiente, Parque Nacional de Guarajonay, La Gomera (reg. number 106051-17099/2014), Consejeria de Medio Ambiente, Cabildo de Gran Canaria (reg. number 5480/2014), Servicio de Medio Ambiente, Cabildo de La Palma (reg. number 2014001631/2014), Ministerio de Medio Ambiente y Medio Rural y Marino, Parque Nacional de la Caldera de Taburiente (reg. number 269046-REUS 52472/2014), Fuerteventura Reserva de la Biosfera, Cabildo de Fuerteventura (reg. number 2348/2014), and the directors of the Parque Natural da Madeira and the Parque Ecológico do Funchal on Madeira for collecting permits. We are very grateful to our research groups, especially to Ignacio Lucas Lledó, Christian Wurzbacher, and Maribet Gamboa, and two anonymous reviewers for their constructive comments on this work. This is publication number 41 of the Berlin Center for Genomics in Biodiversity Research. This work was supported by the Leibniz Association (PAKT für Forschung und Innovation) project FREDIE (SAW-2011-ZFMK-3 to M.T.M.) and by a travel award from the Leibniz-Institute of Freshwater Ecology and Inland Fisheries to S.R.. Individual support was provided by the Swiss National Science Foundation (Early PostDoc.Mobility fellowship P2SKP3_158698 to S.R.), the Japan Society for the Promotion of Science (Long-Term Research Fellowship L-15543 to M.T.M.), the European Investment Funds by FEDER/COMPETE/POCI - Operacional Competitiveness and Internationalisation Programme (POCI-01-0145-FEDER-006958 to S.J.H.), and the National Funds by FCT - Portuguese Foundation for Science and Technology (UID/AGR/04033/2013 to S.J.H. and SFRH/BPD/99461/2014 to P.M.R.).

